# HIV-1-envelope trimer transitions from prefusion-closed to CD4-bound-open conformations through an occluded-intermediate state

**DOI:** 10.1101/2024.07.15.603531

**Authors:** Myungjin Lee, Maolin Lu, Baoshan Zhang, Tongqing Zhou, Revansiddha Katte, Yang Han, Reda Rawi, Peter D. Kwong

## Abstract

HIV-1 infection is initiated by the interaction between the gp120 subunit in the envelope (Env) trimer and the cellular receptor CD4 on host cells. This interaction induces substantial structural rearrangement of the Env trimer. Currently, static structural information for prefusion-closed trimers, CD4-bound prefusion-open trimers, and various antibody-bound trimers is available. However, dynamic features between these static states (e.g., transition structures) are not well understood. Here, we investigate the full transition pathway of a site specifically glycosylated Env trimer between prefusion-closed and CD4-bound-open conformations by collective molecular dynamics and single-molecule Förster resonance energy transfer (smFRET). Our investigations reveal and confirm important features of the transition pathway, including movement of variable loops to generate a glycan hole at the trimer apex and formation or rearrangements of α-helices and β-strands. Notably, by comparing the transition pathway to known Env-structures, we uncover evidence for a transition intermediate, with four antibodies, Ab1303, Ab1573, b12, and DH851.3, recognizing this intermediate. Each of these four antibodies induce population shifts of Env to occupy a newly observed smFRET state: the “occluded-intermediate” state. We propose this occluded-intermediate state to be both a prevalent state of Env and an on-path conformation between prefusion-closed and CD4-bound-open states, previously overlooked in smFRET analyses.

## Introduction

The human immunodeficiency virus type 1 (HIV-1) envelope glycoprotein (Env), composed of gp120 and gp41 subunits that form a homotrimer, is a marvel of viral engineering, able to merge viral and target-host cell membranes, while resisting neutralization by most Env-directed antibodies^1^. Env-facilitated entry of HIV-1 utilizes a double-lock mechanism^2^ comprising initial recognition by a prefusion-closed Env trimer of the primary cellular receptor, CD4, which induces structural rearrangements into a prefusion-open trimer recognized by a co-receptor (CXCR4 or CCR5). Co-receptor recognition induces the formation of a pre-hairpin intermediate spanning viral and cellular membrane, which resolves into a highly stable 6-helix bundle (the postfusion conformation of Env), inducing membrane merger^3^.

Substantial characterization of Env entry has been made including the determination of residue-level static structures of the prefusion-closed Env trimer^4^, various CD4-bound Env trimers^5–8^, the CD4-bound, co-receptor-bound Env^9^, and various antibody-bound states^8, 10–12^. Recent results from EM-tomograms of viral Env with CD4-VLP provide a context for these structures revealing initial contact to occur between a single-extended CD4 and Env trimer in its prefusion-closed conformation^13, 14^ when viral membrane and target membranes are ∼150 Å apart; more extensive contacts and rearrangement of two or three CD4-binding protomers bound per Env trimer are observed when membranes are ∼120 Å apart^15^.

Single-molecule Förster resonance energy transfer (smFRET) has provided information on the prevalence and transition between Env states on virus^16–18^. Three prevalent prefusion-Env states have been characterized, which correspond to (I) a pretriggered ground state that remains to be characterized at the residue-level^18^, (II) a prefusion-closed state that is a requisite transition intermediate and corresponds to the prefusion-closed conformation of Env trimer observed in most ligand-free Env trimer crystal and cryo-EM structures^10, 19, 20^, and (III) an open state that is recognized by CD4 and antibodies that neutralize only laboratory-adapted strains of HIV^5, 16^.

While static structural details of and transition dynamics between prefusion-closed and CD4-bound conformations of HIV-1 Env have been described, dynamic features between these two structures, specifically for transition pathway structures of HIV-1 Env trimer, remain largely unknown. While attempts have been made to simulate transition conformations, these studies have been limited thus far to only the gp120 subunit and have not included full glycans^21^.

Typically, large conformational changes of protein structure occur on timescales ranging from several milliseconds to seconds^22^. Despite substantial advances in computational power^23^, conducting conventional molecular dynamics simulations for proteins on a macroscopic scale is still not feasible. While adopting a coarse-grained approach can simplify the parameterization of protein and glycan residues and enhance computational speed^24^, it is often essential to capture and comprehend the detailed intricacies of an all-atom system.

To tackle the challenge of large-scale simulation of HIV-1 Env trimer, here we employed collective molecular dynamics (coMD) simulation^25^. This approach allowed us to sample conformational change on a macroscopic scale with a reasonable computational timeframe while preserving the complete set of atoms within an all-atom system. We focused on the dynamic process of the HIV-1 Env trimer as it transitions from its prefusion-closed conformation (state II) to its CD4-bound open conformation (state III), using molecular dynamics to identify key characteristics of the transition process. Notably, through our investigation into the transition trajectory, we identify a prevalent intermediary phase, which we name the “occluded-intermediate” phase, and further use smFRET to provide insights into this distinct state, which our results indicate to be a prevalent state, previously overlooked in smFRET analyses.

## RESULTS

### Collective molecular dynamics simulation effectively simulates large conformational changes between two known structures

The coMD simulation necessitates two established structures that serve as endpoints. These endpoints were prepared by modeling homologous template structures, which were used as input files for the coMD simulation. The detailed procedure for the coMD algorithm is provided in Methods section. The outcome of the simulation trajectory was subsequently used for further analyses such as characteristics, glycan coverage and root mean square deviation (RMSD) (Fig. 1a).

**Fig. 1:**
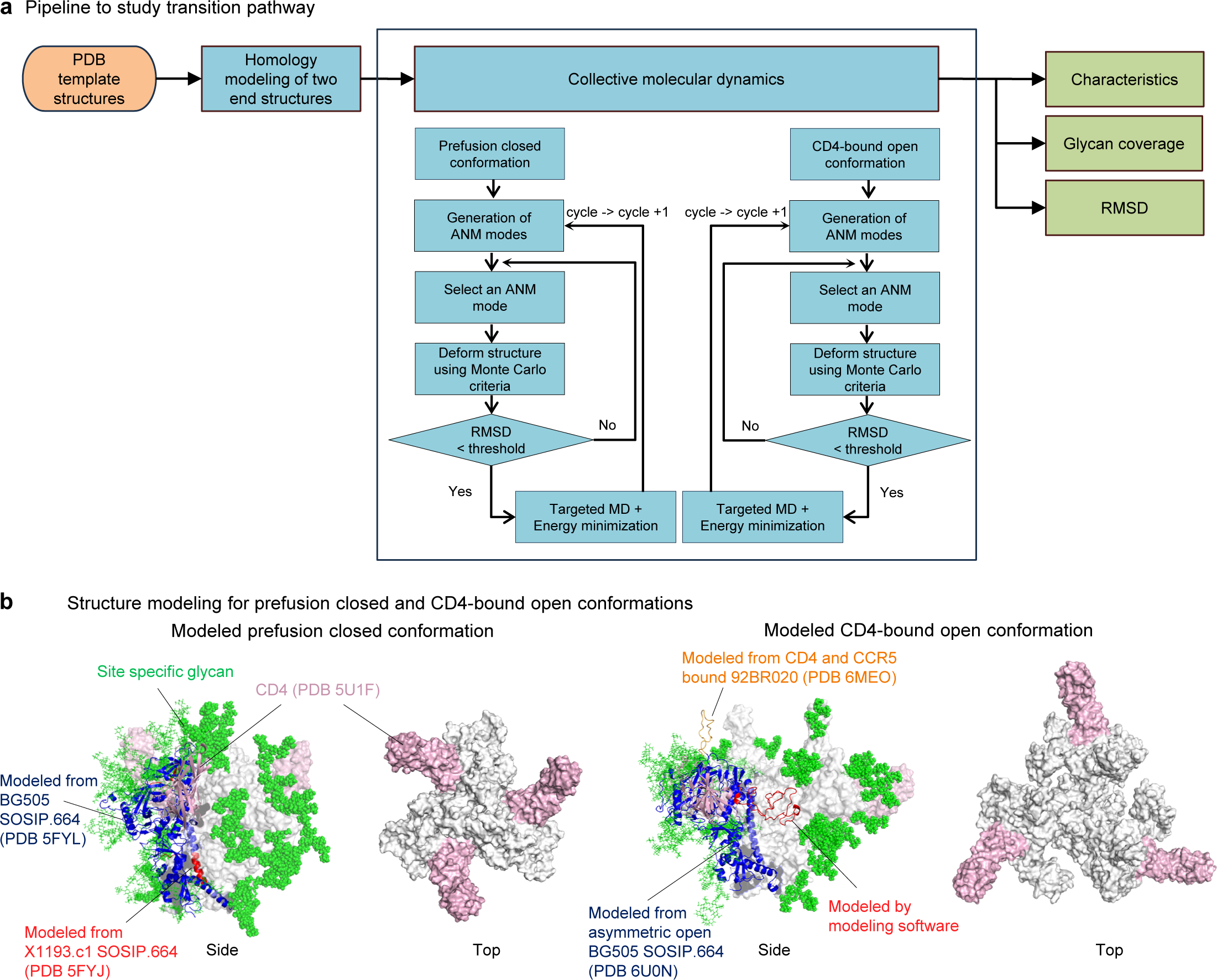
HIV-1 Env transition pathway between prefusion closed to CD4-bound open starts from site specific glycosylated two modeled structures. **a** Pipeline to simulate transition pathway. Input was represented as orange circle and analyses are in blue squares. Pipeline output for homology modeling is shown in **b**, for characteristics in Fig. 2, for glycan coverage in Fig. 3, and for RMSD in Fig. 4. *ANM = anisotropic network model, *RMSD = root mean square deviation **b** Schematic figures of homology modeled prefusion closed and CD4-bound conformations viewing from side and top. Glycans are not shown in top view for clear view. CD4s are in pink, site specific glycans are in green. For prefusion closed conformation, BG505 SOSIP.664 colored in blue was used for main template along with X1193.c1 SOSIP.664 in red. (Left two) In CD4-bound open conformation, BG505 SOSIP.664 asymmetric open in blue was used for main modeling template and V3 loop was adopted from CD4 and CCR5 bound HIV-1 trimer. V1V2 loop were relied on homology modeling software, YASARA (http://www.yasara.org) (Right two).

To understand the HIV-1 Env trimer transition pathway, we set the two initial endpoints structures as prefusion closed and CD4-bound open conformations. We chose to use Env trimers of the BG505 strain^26^ from clade A, as it is one of the extensively studied strains. Due to the constrained dynamic motion by stabilizing structures involving disulfide bond (DS, SOS) or mutation of isoleucine to proline (IP), the study used wildtype (WT) that provides inherent flexibility. The homology modeling for prefusion-closed structure was performed to construct the BG505 WT first endpoint model of Env trimer with BG505 SOSIP.664 (ref ^27^; PDB ID 5FYL) and X1193.c1 SOSIP.664 (ref ^27^; PDB ID 5FYJ) as templates.

For CD4-bound open conformation, the strain was also BG505 WT and two templates, BG505 SOSIP.664 (ref. ^7^; PDB ID 6U0N) and CD4 and CCR5 bound HIV-1 trimer (ref ^9^; and PDB ID 6MEO), were used to construct second endpoint of the simulation. Remaining missing loops were generated by homology modeling software (http://www.yasara.org)^28^. In both endpoint structures, three CD4s were added to allow investigation into the CD4-triggered opening of the Env trimer. To finalize the modeling process, we built site-specific glycans to the entire protein surface following completion of protein modeling (Fig. 1b).

Two simulations were performed in parallel, starting from two endpoints - prefusion closed and CD4-bound open structure, with the objective of reaching the opposite side of each endpoint. Subsequently, the two completed trajectories were merged into a single overall trajectory that encompassed the entire pathway from prefusion closed to CD4-bound open state.

### The transition from prefusion-closed to CD4-bound open conformations reveals essential characteristics of Env trimer opening

The single overall trajectory illustrates the pathway of trimer opening, notably showing that V1V2 and V3 loops that covered the apex of the prefusion-closed conformation to rearrange as it begins to transition to the open state (Fig. 2a). Specifically, the opening process of gp120 involves more than simply expanding the space in the central core; instead, it involves a twisting movement of the V1V2 and V3 loops, causing them to rotate outward. At 50% of the transition pathway, the V1V2 loop becomes disordered, while the V3 loop remains on top of gp120. When the trimer is completely opened, the V1V2 loop moves to the side of Env and V3 loop extends vertically away from the viral membrane (Fig. 2b and Movie S1). The HR1 (HR1_C_) helix, located at the C-terminus of gp41, is initially a disordered loop in the prefusion-closed conformation and starts to assemble, with a distinct helical extension evident at the 50% opening stage – and as the trimer opens further –2.5 to 3 turns in the helix are observed (Fig. 2c). In addition to HR1_C_ extension, α0 at the gp120 behaves in a similar manner by generating helical coil (Fig. 2d). The beta strands β2 and β3, which form a part of a quartet of strands (β20, β21, β2, β3) enclosed by the V1V2 and V3 loops in the closed conformation undergo a flipping motion upon opening (Fig. 2e) to form the 4-stranding bridging sheet. The fusion peptide at the N-terminus of gp41 is initially folded and situated alongside gp41 in the closed conformation. As the transition starts, the loop begins to transition into a coiled coil helix, elongating towards the central core of the trimer (Fig. 2f). All numerical sequences are in HXB2 format, and the corresponding sequence can be found in Supplementary Fig. 1. Overall pathway of transition is shown in Movie S2.

**Fig. 2:**
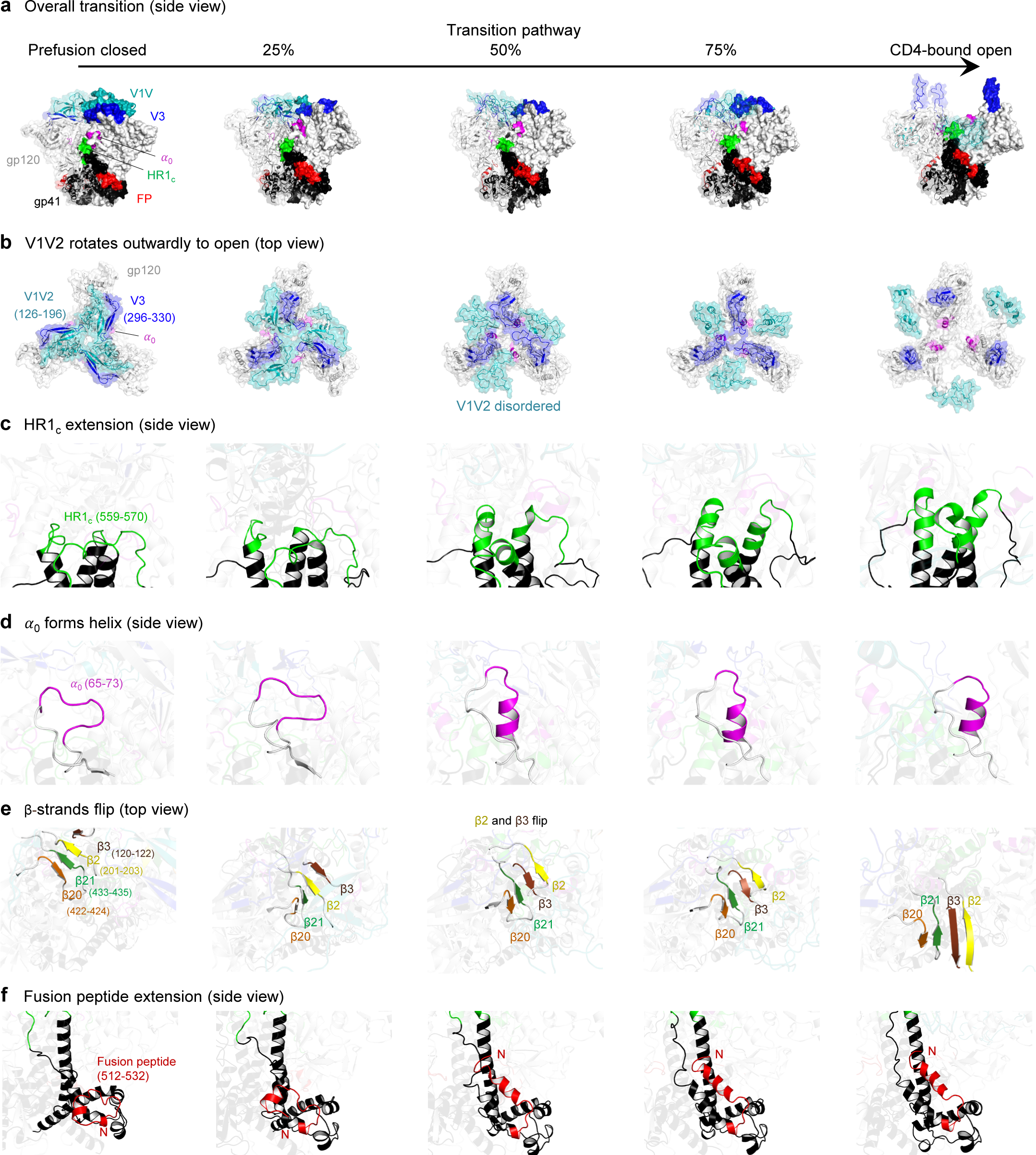
The transition pathway of the HIV-1 Env reveals key attributes of its dynamic opening process. **a** Overall transition pathway from prefusion closed to CD4-bound open conformations viewed from side with distinctive features highlighted using different colors for each characteristic. **b** Three attributes in gp120 v1v2, v3 loops and 𝛼𝛼_0_ are highlighted across the transition pathway viewing from top. **c** The N terminus of gp41, HR1c residues 559-570, are highlighted in green. **d** The 𝛼𝛼0 residues 65-73 are colored in magenta. **e** Four beta strands β20 (orange) residues 422-424, β21 (green) 433-435, β2 (yellow) 201-203, β3 (brown) 120-122 are shown. **f** The fusion peptide residues 512-532 are colored in red and N terminus at 512 is labeled with capital N.

### A large glycan hole in the core of the HIV-1 Env trimer is generated during the transition process

We also investigated glycan coverage of the Env trimer over the transition process using GLYCO^29^. Notably, the V1V2 and V3 loops were extensively covered by glycans throughout the transition. However, a significant glycan hole started to appear being at ∼25% of the opening stage. In the fully opened conformation, the central core of the trimer becomes the most exposed region (Fig. 3a). The glycan hole size was quantified along with the CD4-binding site and base residues. While the size of the glycan hole remained relatively constant at the CD4-binding site and base residues throughout the pathway, there was a noticeable increase in the glycan hole size at the apex when the trimer is fully open (Fig. 3b).

**Fig. 3:**
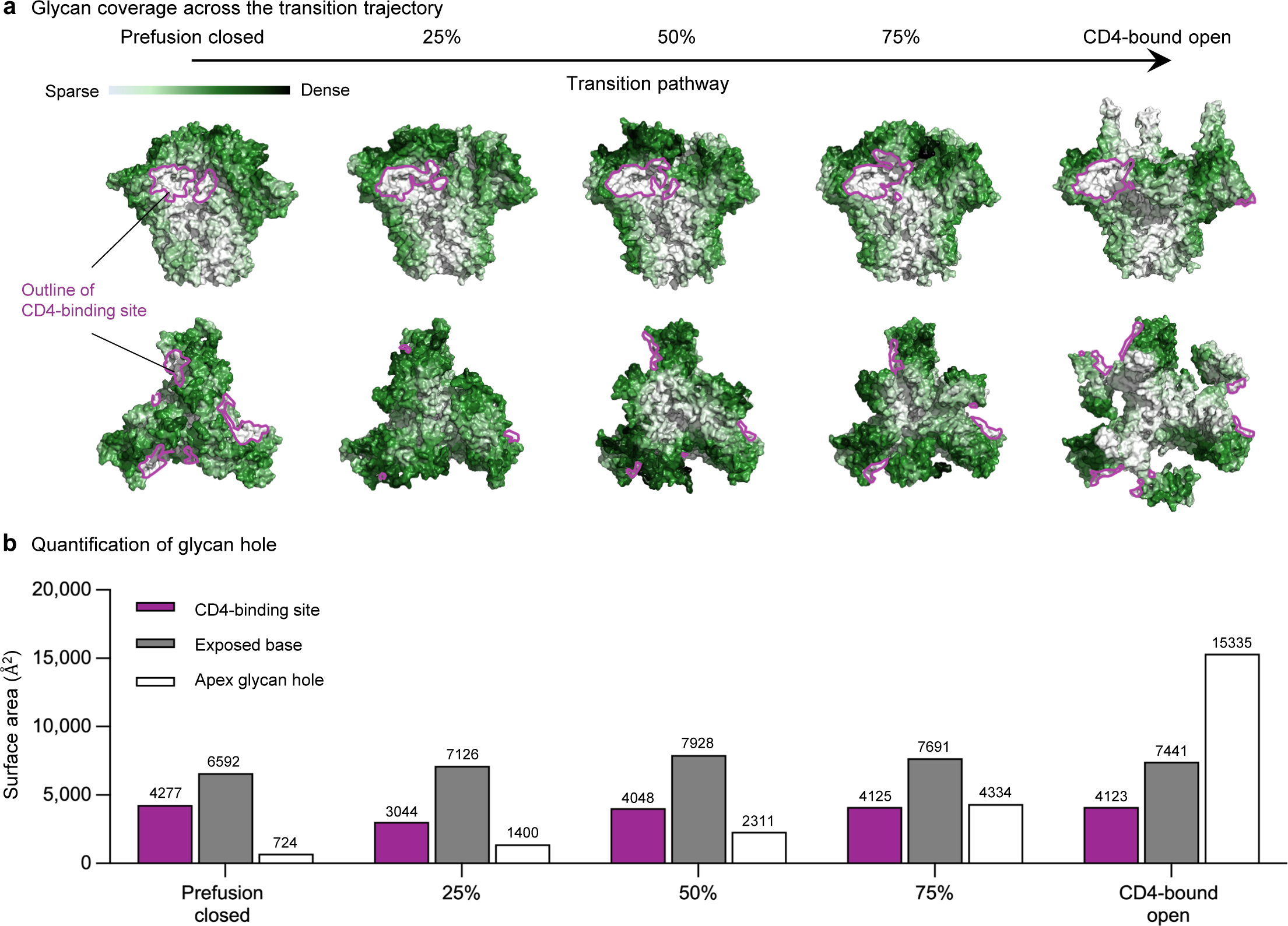
Glycan coverage indicated that a large glycan hole generated during transition pathway. **a** Glycan coverage is colored from white to dark green, while CD4 bound region is represented as dark pink outlines. **b** Quantification of glycan hole size of the top.

### Transition between prefusion closed and CD4-bound states reveals a substantial transition intermediate – the occluded-open conformation

By extracting coordinates from the trajectory, we superimposed all Env trimers bound to antibodies published in Protein Data Bank (PDB)^30^ to examine systematically the structural conformation by RMSD across the transition pathway. The majority of Env trimer in the PDBs had low RMSDs to the prefusion-closed conformation, with 162 PDBs aligning to this phase of the transition process. The CD4-bound open conformation was the second most prevalent conformation, with 14 PDBs aligning to this phase. Notably, our analysis revealed four antibodies – Ab1303^12^, Ab1573^12^, b12^5^, and DH851.3^31^ – that aligned with the intermediate phase of the transition (from 17% to 83% of the transition). The conformation recognized by these antibodies has been described as “occluded” by Ward and colleagues^5^ and as “occluded open” by Bjorkman and colleagues^12^; we have named this conformation the “occluded intermediate” conformation, because of its prevalence in the transition pathway (Fig. 4a).

**Fig. 4:**
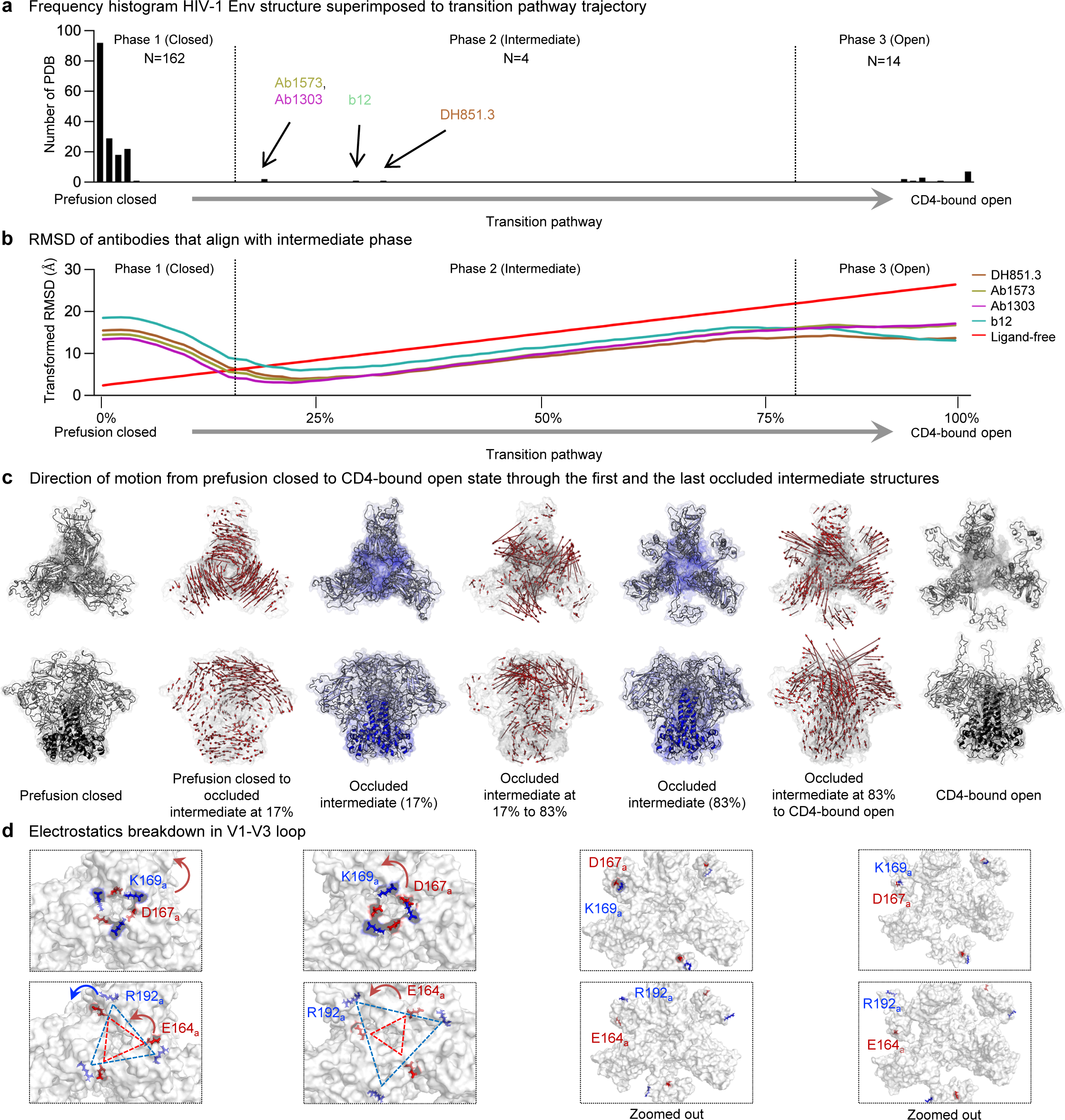
The majority of the transition trajectory demonstrates occluded intermediate conformations. **a** A histogram illustrating the count of PDB structures (N = 180) of HIV-1 Env trimers superimposed onto the trajectory of the transition pathway. Antibodies induce occluded intermediate conformation are highlighted with distinct colors. **b** The RMSDs of antibodies that were identified as inducers of the occluded intermediate state. The RMSD values are linearly transformed based on the RMSD of the prefusion closed conformation (Ligand-free) for comparison. **c** The direction of motion vectors is represented by red arrows from prefusion closed to the initial (17% opening), to the last (83% opening) occluded intermediate and to the CD4-bound open conformation in the trajectory. **d** The major electrostatic interactions in V1-V3 loop in gp120 are depicted in blue for positively charged residues and red for negatively charged residues across the transition pathway. The chain IDs are denoted as subscripts.

The RMSD analysis between the four occluded intermediate conformations and the transition trajectory reveals that a substantial portion of the opening process occurs within the occluded intermediate phase (phase 2, Fig 4b). To delineate conformational changes within the trajectory, we analyzed the direction of motion of each residue as they transition from the prefusion closed to the CD4-bound open, through the occluded intermediate conformation. The major difference of occluded intermediate form comes from V1-V3 loop in gp120, while gp41 remains largely unchanged throughout. The significant movement in gp120 is observed in phases 1 and 3, transitioning from fully closed and to open conformation, in contrast, only local rearrangement occurs in the occluded-intermediate phase (phase 2) (Fig. 4c). In our effort to examine gp120 more closely, we observed that the inter-chain electrostatic interactions (K169 – D167 and R192 – E164) which stabilize the prefusion closed conformations start to weaken as the structure undergoes the process of opening (Fig. 4d). Interestingly these charged residues, particularly D167 and R192 are conserved in sequence (ref), highlighting the importance of these particular features to Env conformational motions.

### Occluded-intermediate conformation can be induced by antibody

Three of the antibodies responsible for occluded intermediate conformations (Ab1303, Ab1573, and b12) target the CD4-binding site, while antibody DH851.3 targets four glycans (N241, N289, N334, and N448) on silence face of Env trimer (Fig. 5a). To investigate the differences between occluded intermediate conformations in both prefusion-closed and CD4-bound open conformations, we visualized the direction of motion using vector formatting. Similar to what we observed in the occluded intermediate state within the trajectory in Fig. 4c, the occluded intermediate conformation showed notable distinctions primarily in the gp120 region when compared to the prefusion-closed conformation. However, the overall direction of motion between the occluded intermediate and CD4-bound open conformations was minimal, except for slight alterations in the gp120 region. This observation implies a remarkable similarity between the occluded intermediate conformations triggered by antibodies and the CD4-bound conformation in phase 2 (Fig. 5b).

**Fig. 5:**
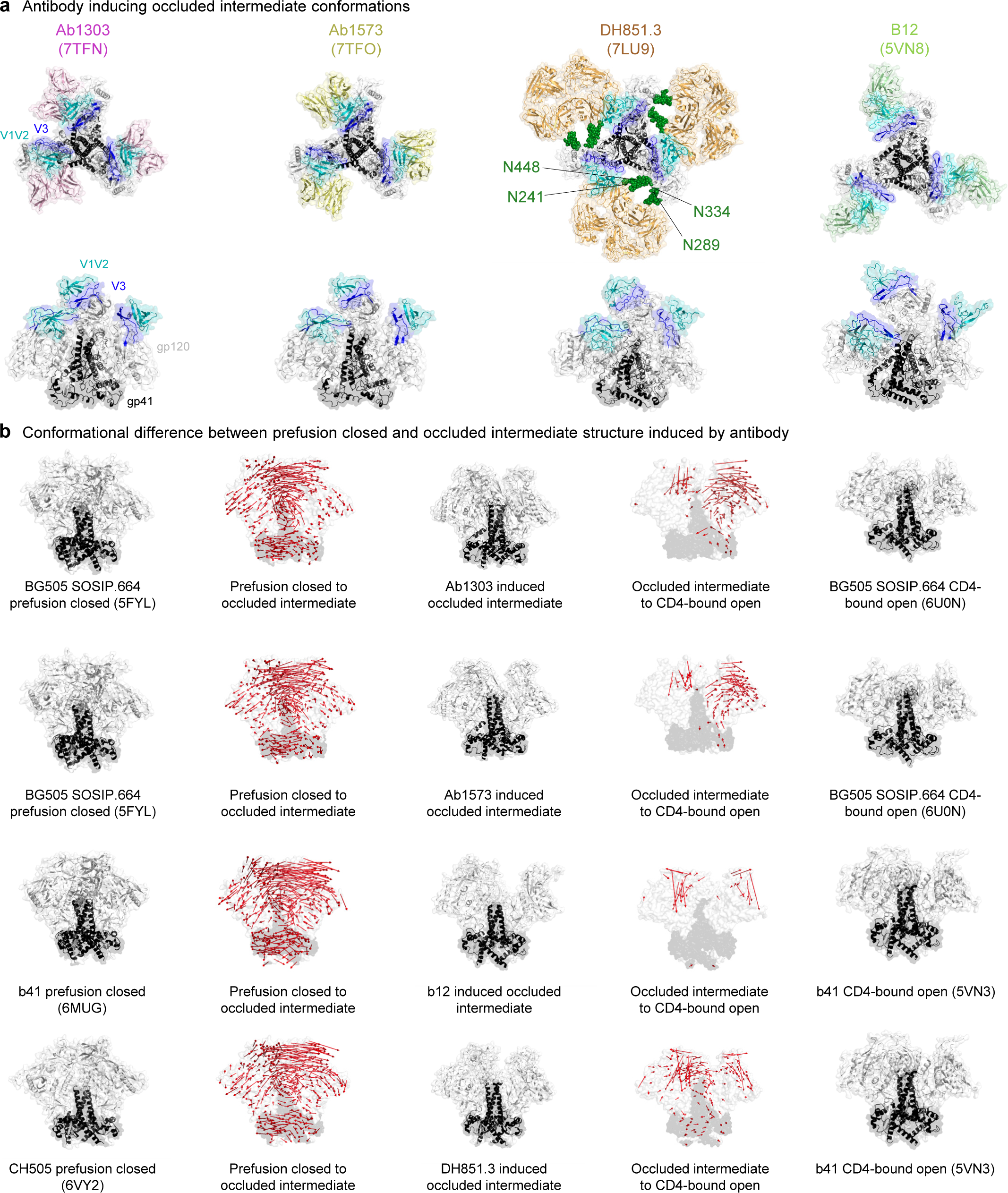
Occluded intermediate conformation can be induced by an antibody. **a** Four antibodies that induce the occluded intermediate conformations. PDB code is enclosed within parenthesis. **b** The direction of motion vectors are indicated by red arrows from the prefusion closed to the antibody induced occluded intermediate and to the CD4-bound open conformation. In the CD4-bound open conformations, the V1V2 and V3 loops are largely absent, leading to missing vectors in that region between the occluded intermediate to the CD4-bound open. Since the CD4-bound open for CH505 is not determined, b41 was used as reference structure. PDB code is shown in parenthesis.

### smFRET reveals a prevalent state of Env on native virions to be stabilized by antibodies that recognize the occluded-intermediate conformation

We utilized smFRET analyses of Env trimer in the context of intact virions (Fig. 6a and Methods) to investigate whether antibodies that recognize the occluded-intermediate conformation revealed in MD simulation could induce conformational reorganization of Env on intact virions. We tested the effect of Ab1303, Ab1573, and DH851.3 on BG505 Env (Env_BG505_) and b12 on Env_JR-FL_ because b12 does not neutralize BG505. Consistent with previous smFRET observations^16–18^, the FRET histogram–conformational population revealed that ligand-free native Env samples three primary states, including the predominant pre-triggered (PT), the prefusion closed (PC), and the CD4-bound open (CBO) (Fig. 6b). In comparison to the ligand-free Env_BG505_, Ab1303, Ab1573, or DH851.3 induces a similar trend of shifts in conformational landscapes of Env_BG505_ and stabilizes a population at mean ∼0.4-FRET that surpasses the one with mean ∼0.3-FRET (CD4-bound open state) (Figs. 6c - e). Similarly, the conformational propensity of Env_JR-FL_ was also measurably shifted in the presence of b12 towards a prevalent state populated at mean ∼0.4-FRET (Figs. 6f, g). A population shift was also observed using blind/unbiased population contour plots complied of FRET trajectories over the first ten seconds (Supplementary Fig. 2). Statistically speaking, three-state Gaussian model fitting of FRET histograms that include a previously well-characterized CBO (∼0.3-FRET) was unfavorable (Supplementary Figs. 3a - b) than that includes a newly populated state (∼0.4-FRET) in Fig.6. Collectively, each of the four antibodies identified in the MD simulation causes conformational shifts of Env and stabilizes Env to favor a ∼0.4-FRET populated state in a similar manner. Therefore, we tentatively assigned the ∼0.4-FRET populated state to the OI state in multi-state Gaussian models and then justified by the following observations and reasoning. While it is difficult to distinguish two different conformational states populated at adjacent FRET levels (∼0.3-FRET vs. ∼0.4-FRET) in the four-state Gaussian model, statistic comparisons between smFRET results of newly identified antibodies (Supplementary Fig.S3) and previously reported smFRET data of several other antibodies (Supplementary Fig.S4) gave us some clues. Previous smFRET studies^16–18^ have revealed stabilization of a prior state sampled by ligand-free Env upon interacting with different classes of antibodies or ligands. Revisiting and refitting previously reported smFRET data using a 4-state Gaussian model were provided in Supplementary Figs. 4c-g and fitting parameters were listed in Figs. 4h. One representative four-state FRET trajectory of virus Env_BG505_ was shown in supplementary Fig.S4g. These new fits (Figs.4c-g) and observed four states in FRET trajectories (an example in Fig.S4g) suggest the existence of the new OI state (∼0.4-FRET) in the previous datasets. Interestingly, the population of the OI state was not enriched in the presence of any of the previously tested antibodies. This observation contrasts with the results in Supplementary Fig.S3c, in which the OI state was measurably enriched with antibodies Ab1303, Ab1573, DH851.3, or b12. These antibodies were the same antibodies identified in the computational analysis (Fig.4a). Thus, the consistent enrichment of a ∼0.4-FRET population of Env induced by Ab1303, Ab1573, DH851.3, or b12 implies that this ∼0.4-FRET population may be a previously uncharacterized prior conformation of Env distinct from CD4-bound open. This distinct conformation is likely the occluded-intermediate state identified by MD simulations. Indeed, distances between the donor and acceptor labeled on Env in a three-phase transition path from pre-fusion closed to occluded intermediate to CD4-bound open are in good agreement with dominant FRET levels in histograms (Supplementary Fig. 5). Therefore, we reason that the occluded intermediate (OI) is the 0.4-FRET state, which is likely buried within the CD4-bound open state in the three-state model and prevails after Env binds to a specific class of antibodies.

**Fig. 6:**
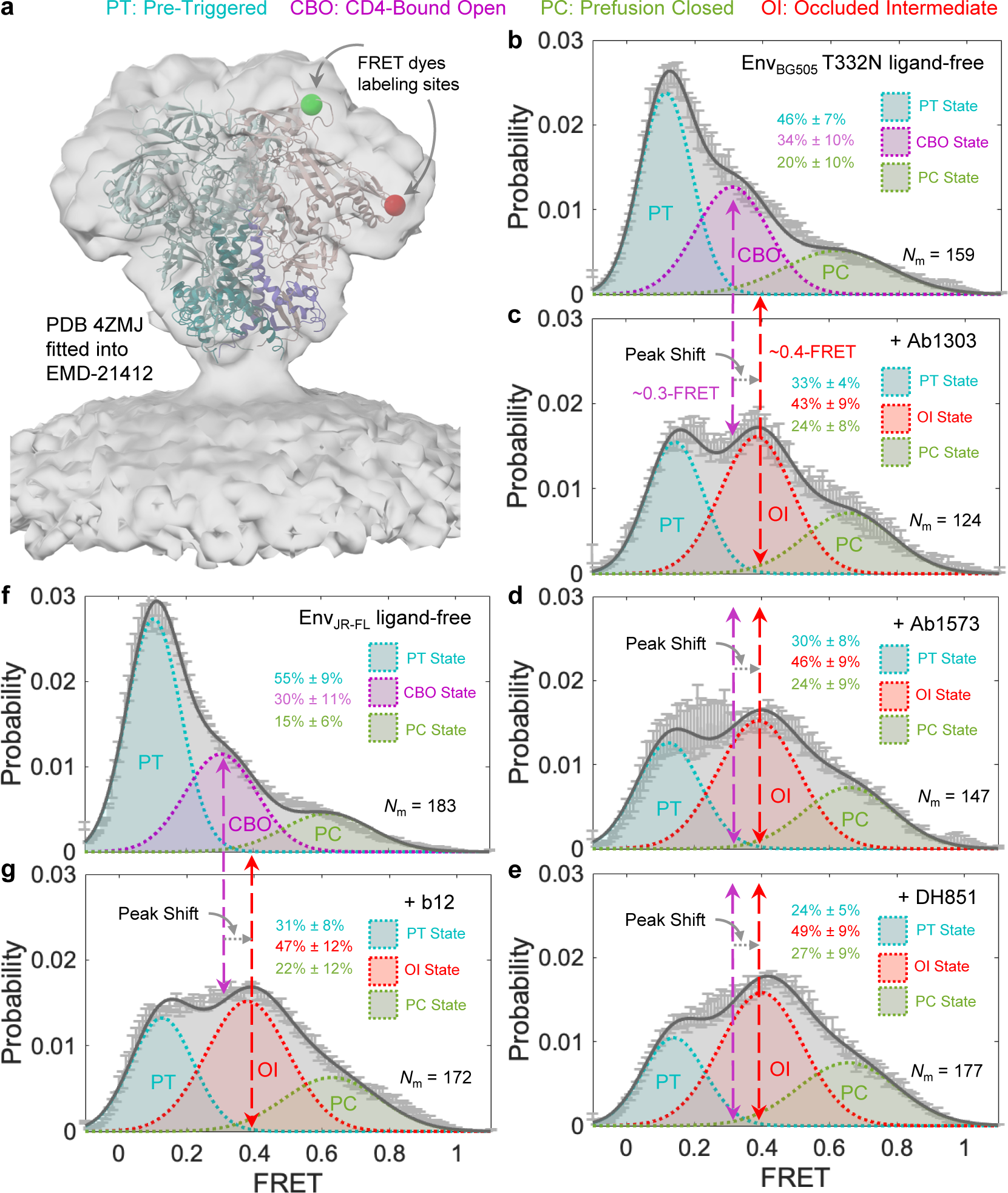
Antibodies induce and stabilize Env to sample a conformation distinct from the CD4-bound open conformation revealed by smFRET imaging of full-length Env on native virions. **a** Demonstration of fluorescent donor-acceptor labeling sites on an Env_BG505_ trimer^22^ in the context of native virions used in smFRET studies. The structure of the Env trimer (PDB access code 4ZMJ)^24^ was fitted into the electron density map of the membrane-present Env trimer (access code EMD-21412)^15^. Two fluorophores were site-specifically labeled to two sites (green ball – donor labeling site, red ball – donor labeling site) located in variable loops V1 and V4, respectively, of a single protomer (gp120 monomer in light pink and gp41 in magenta) within an Env trimer (wildtype protomers in light cyan/dark blue). **b** FRET histogram, indicative of conformational ensemble of ligand-free Env_BG505_ T332N present on native virions. Ligand-free Env_BG505_ predominately occupies the pre-triggered (PT) conformation among three primary conformational states (PT, CBO: CD4-bound open, and PC: pre-fusion closed). *N*_m_ number of Env molecules was compiled into the histogram and fitted into a sum of three Gaussian distributions, as previously described^21–23^. **a -e** FRET histograms of Env_BG505_ in the presence of Ab1303 (**c**), Ab1573 (**d**), and DH851 (**e**), respectively. These antibodies induce a similar trend of peak shifting in the Env_BG505_ conformational ensemble and stabilize a conformation (Occluded Intermediate – OI populated at ∼0.4-FRET), distinct from the CD4-bound open (CBO populated at ∼0.3-FRET) conformation. Peak shifting was also observed using blind/unbiased population contour plots (see Supplementary Fig. 2). Constrained model-fitting of histograms to include a CBO-populated Gaussian was statistically unfavorable (see Supplementary Fig. 2). **f, g** Experiments as in b, FRET histograms of Env_JR-FL_ in the absence (**f**) and presence of b12 (**g**), respectively. Antibody b12 induced a similar peak shift from CBO to OI in the conformational population that Env_JR-FL_ samples, as observed in **c**-**e**.

### New transition-pathway mechanism with occluded intermediate state and the essential common core of each state

Dynamic studies of Env^16–18^ have previously identified three primary states for the Env trimer – pre-triggered, pre-fusion closed, and CD4-bound open (Fig. 7a). The pre-triggered state must transition through an obligatory intermediate, known as pre-fusion closed, to CD4-bound open (which is also known, and the state recognized by CD4-induced antibodies such as 17b and 48D, which recognize the four-stranded bridging sheet). In our current study, we disentangled an additional transitional prefusion state populated at 0.4-FRET (occluded intermediate state) on the fusion pathway. The occupancy of this state can be enriched by a specific class of antibodies. coMD transitional-path simulation of a fully glycosylated Env trimer from prefusion closed to CD4-bound open identified distinguishable but conserved structural features across four Ab-bound Env structures, which implies the presence of an occluded intermediate state on the path (Fig. 7a).

**Fig. 7:**
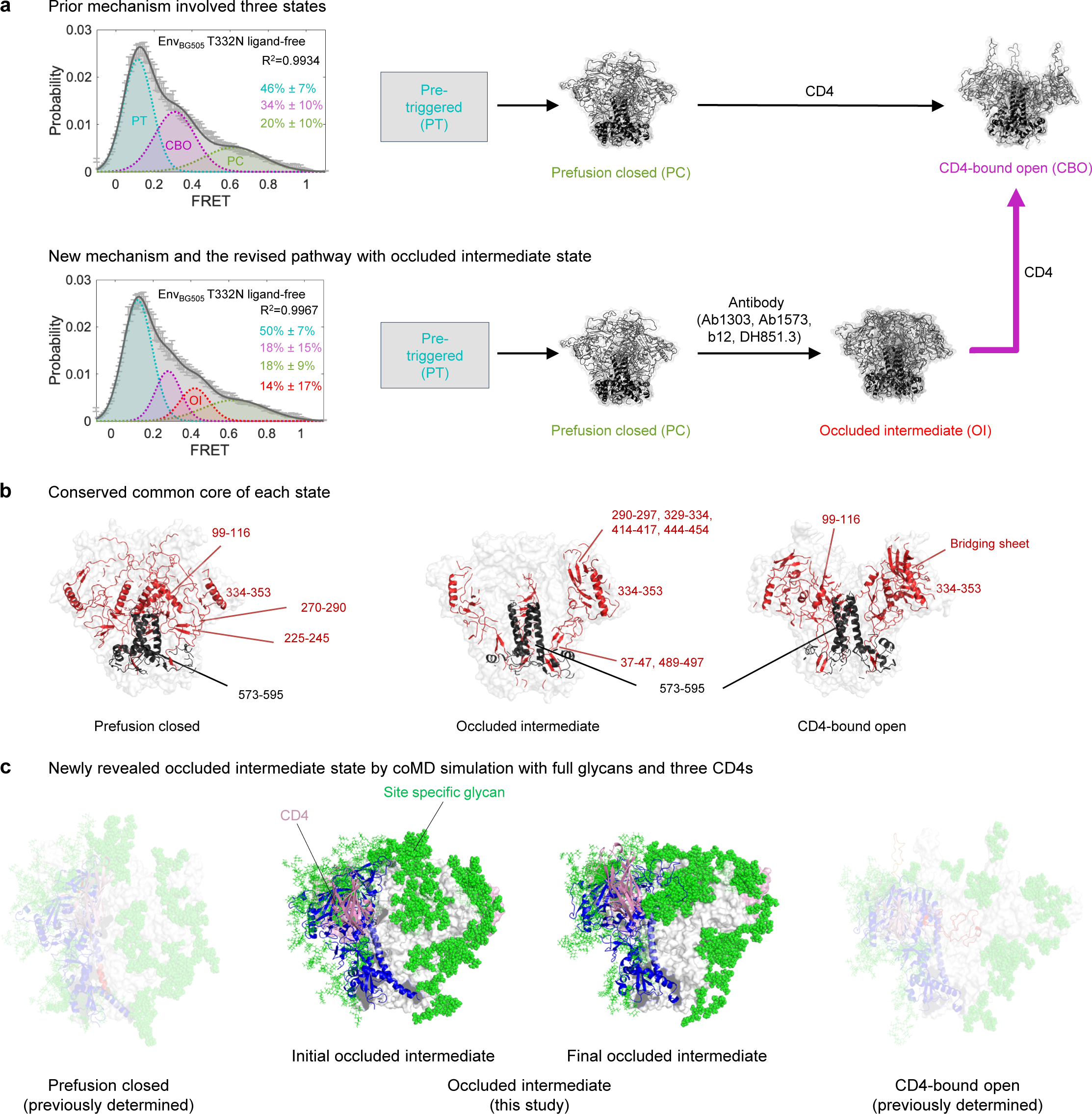
Essential common core of each state and new discovery of the pathway. **a** Prior smFRET analysis and pathway with three states. The pre-triggered state is shown as gray box since its structural information remains unidentified. **b** New mechanism with smFRET and the diagram describes the identification of a new state, occluded-intermediate. The new discovery from the coMD simulation is denoted by magenta arrow. **c** The conserved core of each state analyzed by pairwise distances of less than 5 Å. **d** Initial and final state of occluded intermediate structure with full glycans and CD4 bound. Prefusion closed and CD4-bound open conformations are also shown but faint.

To identify the difference between three states (prefusion closed, occluded intermediate, and CD4-bound open), we analyzed the conserved structural core of each state by superimposing PDB structures within those respective states. Through this approach, we were able to examine the preserved characteristics present across these structures (Fig.7b). The most prominent feature between three states was the conservation of the gp41 core (residue 573-596), except for the fusion peptide and HR1_C_ region in three states. This provides the evidence that conformational dynamics occurs mostly through gp120 (Fig. 7b).

In the context of the prefusion closed state of gp120, a substantial portion of the V1V2 region was structural preserved along with an alpha helix at residues 99 to 116. In addition, two beta sheets within the residue ranges of 225-245 and 270-290, and a helix at 334-353 are also notably maintained.

The interesting feature of common core for the occluded-intermediate state is the beta sheet formed by N- and C-terminus of gp120 is preserved (residue 37-47 and 489-497). Furthermore, beta sheets observed to play a key role in structuring the occluded intermediate state — within residue ranges 290-297, 329-334, 414-417, and 444-454 —show conservation. Such structural conservation was also observed for the alpha helix spanning residues 334 to 353, which also formed part of the common core.

In the CD4-bound open conformation, similar to the occluded-intermediate state, a beta sheet created by the N-terminus and C-terminus of gp120 (residues 37-47 and 489-497) remains structurally conserved. Additionally, four beta strands are preserved in one protomer and a portion of highly conserved alpha helix located at residue 99-116 is also maintained in this state.

Based on the coMD results, we defined occluded intermediate as an intermediate phase of the transition trajectory (Fig.7c). The initial occluded intermediate closely resembles the prefusion closed conformation, while the final occluded intermediate exhibits characteristics more towards the CD4-bound open conformation, with the glycans folded closer to the Env trimer surface (Fig. 7c).

## DISCUSSION

We performed simulations complemented by experiments to explore the transition pathway trajectory of fully glycosylated Env trimers from prefusion closed to CD4-bound open conformations to achieve a better understanding of the underlying entry mechanism. Our results highlight the central role of a previously little-understood structural intermediate – the occluded-intermediate state – within the Env-mediated fusion pathway, which begins with the pre-triggered state and progresses to the formation of post-fusion six-helix bundle (Supplementary Fig. 6). The trajectory obtained in this study is first discovery of full transition pathway with site specifically glycosylated Env structures. This transition pathway of Env can be compared and aligned with any available structures of Env trimers, including newly obtained ones (Fig. 4). This approach resulted in assigning Env structures to well-ordered phases along the transition trajectory and further characterizes phased Env states (Figs.1-5). Recently, cryoET of Env in membranes by Li et al.^15^ unraveled the initial contact of Env with CD4, and cryoEM single particle analysis of engineered Env by Dam et al.^14^ characterized one- or two-CD4-bound Env trimer as asymmetric prefusion-closed or open intermediates, respectively. Notably, our trajectory results indicate that each of the protomers transitions through the intermediate state. Overall, our trajectory approach integrates Env dynamics and static structures, expanding our understanding of the structural features of Env trimers in the context of its transition process upon engagement with CD4 and the host (Supplementary Fig. 6).

Our coMD simulations revealed a segmentation-like three-phase trajectory (Figs. 4a and b), in which transition predominantly occurs via an occluded-intermediate state (Phase 2) in between prefusion closed (Phase 1) and CD4-bound open states (Phase 3). Env structures complexed with an antibody (including ab1303, ab1573, b12, and DH851.3) lay in Phase 2, whereas 162 available prefusion closed structures are in Phase 1, and 14 CD4-bound open structures are in Phase 3. This three-phase trajectory strongly indicates the presence of a previously unknown fourth state that the above-mentioned four antibodies specifically recognize, the occluded-intermediate state, in addition to the well-known pre-triggered, prefusion-closed, and CD4-bound open states (Supplementary Fig. 6).

This prevalence of the occluded intermediate state was further validated through smFRET imaging of virus Env in the presence of each of four specific antibodies, ab1303, ab1573, b12, and DH851.3. smFRET revealed a repetitive pattern of peak shifts towards ∼0.4- FRET in Env conformational distributions when a specific antibody was present. The parameters chosen for model fitting of FRET histograms were determined based on visual inspection of FRET trajectories and idealization using hidden Markov modeling^16–18, 32^. Constrained model-fitting of FRET histograms of tested Ab-bound Env to include a CBO (∼0.3-FRET) was statistically unfavorable (Supplementary Figs. 3a - b) than the one with OI state (∼0.4-FRET) in Fig.6. Model-fitting of Ab-Env FRET histograms with four states increased the likelihood of overfitting and the uncertainty of state occupancy (Supplementary Figs.3c -d), given the adjacency between ∼0.3-FRET and ∼0.4-FRET. We revisited and performed a 4-state Gaussian model fitting of previously published smFRET datasets of several other tested antibodies and revealed inconsistency with regard to the enrichment of the OI state. None of the previously tested Abs was able to increase the population of the OI state to a measurable level (Supplementary Fig.4), whereas all antibodies identified in the MD simulation were able to (Supplementary Fig.3c and Fig.6). At the single-molecule level, the ∼0.4 FRET populated signal in single FRET trajectories can be distinguished and idealized by HMM (Fig.S4g). Therefore, we related this new peak-indicated conformation to the occluded intermediate (OI) state. Env is highly dynamic on the virus surface, likely spontaneously having access to the occluded intermediate state as well as other three states. The 3-state or 4-state model fitting was used to show the prevalent state with statistics. Because the FRET signals of CBO and OI states were so close due to the current labeling sites, the four-state model will be prone to overfitting. The 3-state model that includes CBO does not mean the lack of OI presence. Similarly, the 3-state model that includes OI does not mean the lack of CBO presence. Our strategy using different models is to show whether CBO or OI prevails in different conditions. The reason why this new smFRET state (OI) did not prevail, so it was not previously captured, is multi-fold: 1) in the 2-CD4-bound state, where a third of the protomers are in the intermediate conformation, the smFRET difference between occluded intermediate and CD4-bound open state may have been too small to enable its prior identification; 2) the relative occupancy of Env in this new state may only prevails when specific antibodies are present, and/or 3) the four newly characterized antibodies recognizing this state were not assessed in previous studies. Thus, specific antibodies that preferentially recognize this new state stabilize Env and further facilitate the revealing of this state.

The occluded intermediate is an on-path conformational state of Env during fusion, necessitating its candidacy as a target for developing vaccines, antibody therapy, and inhibitors. The four antibodies, ab1303, ab1573, b12, and DH851.3, that recognize the occluded intermediate state show properties that may be of vaccine relevance; certainly antibody b12 neutralizes tier II-resistant isolates, and antibodies ab1303 and ab1573 were elicited by immunization and exhibit weak neutralization^31, 33^. It will be interesting to see if effective vaccine-elicited neutralizing antibodies will continue to be constrained to antibodies that target the pretriggered or prefusion-closed states, or if they will also be elicited against the occluded-intermediate state.

## METHODS

### Collection of HIV-1 Env trimers and RMSD analysis with the transition trajectory

Approximately 500 PDBs of antibody bound HIV-1 Env complexes were obtained from SAbDab^34^ on March 23^rd^, 2023. From this set of structures, those identified as scaffold or peptide structures were excluded. Initially 178 trimers with complete gp120 and gp41 were collected and one was excluded due to computational modeling^35^ (PDB: 3J70). In addition, ligand free HIV-1 Env trimer^19^ (PDB: 4ZMJ), along with one and two CD4 bound HIV-1 Env trimers^14^ (PDB: 8FYI, 8FYJ) were included, resulting in a total 180 PDBs participating in the analysis. All 180 PDBs were aligned with each time frame of transition pathway and RMSD analysis was performed using PyMOL (www.pymol.org). RMSD relative to trajectory is minimum of all three protomers. The boundary between Phase 1 and 2 was defined as the average cross point with the ligand free plot. The inflection point of the four antibodies was set as the boundary between Phase 2 and 3.

### Homology modeling and glycosylation of starting structures

The initial structure of the trajectory, prefusion closed conformation, was modelled using multi templates PDBs 5FYL and 5FYJ. Three CD4s were modelled using template PDB 5U1F. Final structure of CD4-bound open conformation was modeled with two templates, PDB 6U0N and 6MEO. Site specific glycans on each of the two modeled structures were built using CHARMM-GUI Glycan Reader & Modeler^36^.

### Collective molecular dynamics

Collective molecular dynamics (coMD)^25^ runs two separate jobs in parallel, utilizing both initial and final structures, which are defined as the forward and backward jobs, respectively. With an initial structure, the program generates a complete set of spectrums of the Anisotropic Network Model (ANM) modes and subsequently selects an ANM mode based on eigenvalue spectrum. The selected ANM mode goes into the deformed structure using Monte Carlo/Metropolis scheme. Once the selected mode has been accepted by Monte Carlo/Metropolis criteria, the program calculates the RMSD between the structure before and after the ANM mode selection. If the RMSD falls lower than a threshold, the targeted MD simulation is performed along with energy minimization. Forward run targets to the final structure, while the backward run targes to the initial structures during targeted MD. If the RMSD threshold is not satisfied, then the algorithm goes back to the process of selecting an ANM mode and repeats the same protocol until RMSD meets the criteria. The cycle continues until the RMSD between forward and backward structures reaches a predefined threshold of 0.15 Å. Upon completing the cycles, the combined forward and backward trajectories represent the comprehensive full path of the simulation.

### Glycan coverage calculation and quantification of glycan holes

Glycan coverage of transition pathway was calculated by GLYCO^29^ with glycan cutoff 20 Å, surface area cutoff 0 Å to include all protein residues in the calculation. Quantification of the apex glycan hole was performed by placing a sphere with the centroid in between three K117 and three G312 residues with a radius of average distance between the centroid and these residues. The residues on the sphere were defined as the apex glycan hole residues. CD4 binding site residues were collected by taking the non-zero value of surface area difference between CD4-bound antibody complexes as described in Chuang et al^37^ and Env trimer. Base residues were defined as the residues located at the bottom of Env trimer and exposed by glycan. The exposure of glycan was calculated by GLYCO^29^ with a glycan cutoff 83 Å which corresponds to the number of atoms in mannose-5 excluding hydrogen atoms. The summation of surface area for each site was analyzed using FreeSASA^38^.

### Modeling dyes for tag distance analysis

To mimic two dyes for smFRET analysis, we modeled two dyes in each of trajectory frame. For the first dye, peptide sequence GQQQLG was modeled in between residue V134 and T135 within the V1 loop. For the second dye, the peptide sequence GDSLDMLEWSLM was modeled between T399 and S400. The distance between the third residues on each tag was measured.

### Conserved common region analysis

The 96 closed PDB structures were collected. As for the occluded-intermediate state, four antibody induced occluded intermediate structures (PDB: 7TFN, 7TFO, 5VN8, 7LU9) and additional five occluded intermediate state structures from the trajectory (25%, 35%, 45%, 55% and 65% open) were collected. Regarding CD4-bound open conformation, fully open nine conformations (PDB: 5VN3, 6U0N, 6U0L, 6OPP, 6OPO, 6OPQ, 6OPN, 6X5C, 6X5B). The structures from each state were aligned, and the superimposed residues with pairwise distances of less than 5 Å were gathered to analyze conserved common core specific to each state.

### smFRET analysis of Env trimers on HIV-1 viral particles

The preparation of fluorescently labeled Env trimers incorporated onto pseudotyped lentiviral particles and the following smFRET experiments and analysis are as previously described^16–18, 32^. Fluorescently labelable viruses incorporated with Env_BG505_ were produced by co-transfecting HEK293T cells with a mix plasmid of tag-free wildtype vs. Env-tagged full-length Q23 BG505 clones (reverse transcriptase deleted) containing Env_BG505_ with Q23 as the backbone. Env-tagged Q23 BG505 carries Q3 and A4 peptides in variable loops V1 and V4, respectively^17, 18^. Pseudotyped viruses incorporated with Env_JR-FL_, were produced by co-transfecting HEK293T cells with a Gag-Pol packaging plasmid pCMV delta R8.2 (Addgene_12263) and a diluted mix of full-length wildtype tag-free gp160 plasmid (pCAGGS JR-FL gp160) Env_JR-FL_ and dually peptide-tagged Env_JR-FL_ in which peptides Q3 and A4 were introduced to V1 and V4 of JR-FL gp160, respectively^16, 17^. A ratio of 40:1 (tag-free wildtype vs. Env-tagged plasmids) was used to ensure that statistically, one dually peptide-tagged protomer within a trimer in otherwise unlabeled trimers on a single virion was available for fluorescent labeling. Viruses were harvested after 40 h post-transfection, filtered, and concentrated on top of 15% sucrose by ultracentrifugation at 25,000 rpm for 2 h, followed by resuspension in the 50 mM HEPES buffer containing 10 mM MgCl_2_ and 10 mM CaCl_2_^16–18, 32^. The resuspended viruses were incubated with a Cy3 derivative LD555-cadaverine (LD555-CD, 0.5 μM, donor fluorophore), a Cy5 derivative LD655-coenzyme A (LD655-CoA, 0.5 μM, acceptor fluorophore) in the presence of two enzymes, transglutaminase (0.65 µM)^39^ and AcpS (5 µM)^40^ overnight at room temperature. PEG2000-biotin (0.1 mg/ml) was added to the above reaction and incubated for 30 mins at room temperature. Viruses were then purified to remove excessive dyes and lipids by ultracentrifugation over a 6–18% Optiprep gradient at 40,000 rpm for 1 hour.

smFRET imaging of fluorescently labeled viruses was performed on our customized prism-based total internal reflection fluorescence (prism-TIRF) microscope, as previously described^16–18, 32^. Fluorescently labeled HIV-1 viruses were immobilized on a PEG-passivated biotinylated quartz-coverglass sample chamber placed on the microscope. Immobilized viruses were imaged in the 50 mM pH 7.4 Tris buffer containing 50 mM NaCl, a cocktail of triplet-state quenchers, 2 mM protocatechuic acid (PCA), and 8 nM protocatechuic-3,4-dioxygenase (PCD) at room temperature. Where indicated, viruses were incubated with 0.1 mg/ml b12 antibody for 30 mins before imaging, and the antibody was continuously present during the imaging. The evanescent field generated by the total internal reflection of 532-nm single-wavelength laser excitation (Ventus, Laser Quantum) directed to a prism excites donor fluorophores on viruses. Optical signals from both donor and acceptor fluorophores were collected through a water-immersion 60x Nikon objective (1.27-NA), optically separated by a dichroic filter mounted (Chroma) and filtered by two emission filters (ET590/50, ET690/50, Chroma) mounted inside a MultiCam LS image splitter (Cairn Research). Fluorescence signals were recorded on two synchronized sCMOS cameras (Hamamatsu ORCA-Flash4.0 V3) with 40 milliseconds time resolution for 80 seconds.

All smFRET data was analyzed by a SPARTAN software package^41^ and customized MATLAB-based scripts. Fluorescence time series (trajectories) were extracted from image stacks and used to determine FRET efficiency trajectories based on FRET= I_A_/(I_D_+I_A_), where I_D_ and I_A_ indicate the fluorescence intensity of the donor and acceptor, respectively. FRET efficiency trajectories at the single-molecule level were further derived. Each trajectory records time-correlated structural changes of one Env protomer within an Env trimer on a virion, as previously described and characterized^16–18, 32^. Trajectories (as demonstrated in Fig.S4g) that show a single photobleaching point, sufficient signal-to-noise ratio and fluorophore lifetime, and a negative correlation between two fluorophores were included for further data processing. Manual visualization was then used to ensure that only Env molecules with a single hetero dye (Cy3/Cy5) labeled protomer within an Env trimer on a virion were included. These criteria rule out the cases of multiple labeled protomers in a trimer, multiple labeled Envs on one virion, Env lacking either donor or acceptor or both on a virion, and inactive Env. Fluorescence and FRET trajectories that meet our filter settings and manual inspection were included in compiling FRET histograms (indicative of conformational distributions). Under each experimental condition (ligand absence/presence), more than 100 individual trajectories were included. FRET histograms were presented as mean ± s.e.m. and fitted into a sum of three or four distinct Gaussian/normal *N* (*μ*, *σ*^2^) distributions using the least-squares fitting algorithm in MATLAB. Parameters (*μ*, *σ*) chosen for curve fitting were determined based on visual inspection of all traces that exhibit state-to-state transitions and the idealization of individual traces using multi-state hidden Markov modeling^16–18, 32^. The area under each Gaussian curve was further estimated to represent the relative state occupancy of Env. Previous smFRET studies of Env^16, 17^ have revealed that Env samples three primary conformational states, and upon activation by CD4 molecules, Env transits from a pre-triggered (PT state, ∼0.1-FRET) to the CD4-bound completely open (CBO, 0.3-FRET) through a pre-fusion closed (0.65-FRET). Different neutralizing antibodies have indicated their preference for Env conformations present on the viruses in smFRET analysis^16, 18^. In this study, we used similar approaches to test the conformational preference of four different antibodies (Ab1303, Ab1573, DH851, and b12) for full-length Env on virions and revealed a measurable shift in FRET histograms that was not previously observed.

## DATA AND SOFTWARE AVAILABILITY

## Competing Interests

All authors declare no competing interests.

## Acknowledgments

We deeply appreciate J. Krieger for his invaluable support and guidance with coMD. We thank J. Stuckey for assistance with figures, and members of the Structural Biology Section and Structural Bioinformatics Core, Vaccine Research Center, for discussions and comments on the manuscript. We also thank I-T. Teng for assistance with antibodies. Support for this work was provided by the Vaccine Research Center, an Intramural Division of the National Institute of Allergy and Infectious Diseases (NIAID), NIH, by NIH/NIAID R01 AI181600, NIH/NIGMS R35 GM151169, and by the collaborative development award from Duke Center for HIV Structural Biology (NIH/NIAID U54 AI170752) to M. Lu. This work utilized the computational resources of the NIH HPC Biowulf cluster (http://hpc.nih.gov).

## Author Contributions

M. Lee led the simulation of the transition pathway, analyzed data, and wrote the paper. M. Lu performed smFRET, and B.Z. produced antibodies 1303, 1573, DH851.3, DGNM-f.01. T.Z. provided antibody b12. R.K. and Y.H. contributed virus preparation, imaging acquisition for smFRET data analysis. R.R. provided guidance on the simulation. P.D.K. analyzed data and wrote the paper, with all authors providing revisions.

## Notes

### Competing Interest Statement

The authors have declared no competing interest.

