## Supplemental Material for "HIV-1-envelope trimer transitions from prefusion-closed to CD4-bound-open conformations through an occluded-intermediate state"

**gp120**

31 $\alpha_0$
AENLWVTVYYGVPVWKDAETTLFCASDAKAYETEKHNVWATHACVPTDPNPQEIHLENVTEEFNMWKNNM

101 β3 V1V2
VEQMHTDIISLWDQSLKPCVKLTPLCVTLQCTNVTNNITDDMRGELKNCSFNMTTELRDKKQKVYSLFYR

171 β2
LDVVQINENQGNRSNNSNKEYRLINCNTSAITQACPKVSFEPIPIHYCAPAGFAILKCKDKKFNGTGPCP

241 V3
SVSTVQCTHGIKPVVSTQLLLNGSLAEEEVMIRSENITNNAKNILVQFNTPVQINCTRPNNNTRKSIRIG

313
PGQAFYATGDIIGDIRQAHCTVSKATWNETLGKVVKQLRKHFGNNTIIRFANSSGGDLEVTTHSFNCGGE

382 β0 β21
FFYCNTSGLFNSTWISNTSVQGSNSTGSNDSITLPCRIKQIINMWQRIGQAMYAPPIQGVIRCVSNITGL

453
ILTRDGGSTNSTTETFRPGGGDMRDNWRSELYKYKVVKIEPLGVAPTRAKRRVVG

**gp41**

512 FP $HR1_c$
AVGIGAVFLGFLGAAGSTMGAASMTLTVQARNLLSGIVQQQSNLLRAIEAQQHLLKLTVWGIKQLQARVL

582
AVERYLRDQQLLGIWGCSGKLICTTNVPWNSSWSNRNLSEIWDNMTWLQWDKEISNYTQIIYGLLEESQN

652
QQEKNEQDLLALD

**Supplementary Fig. 1: Characteristics in sequence using the HXB2 format.**

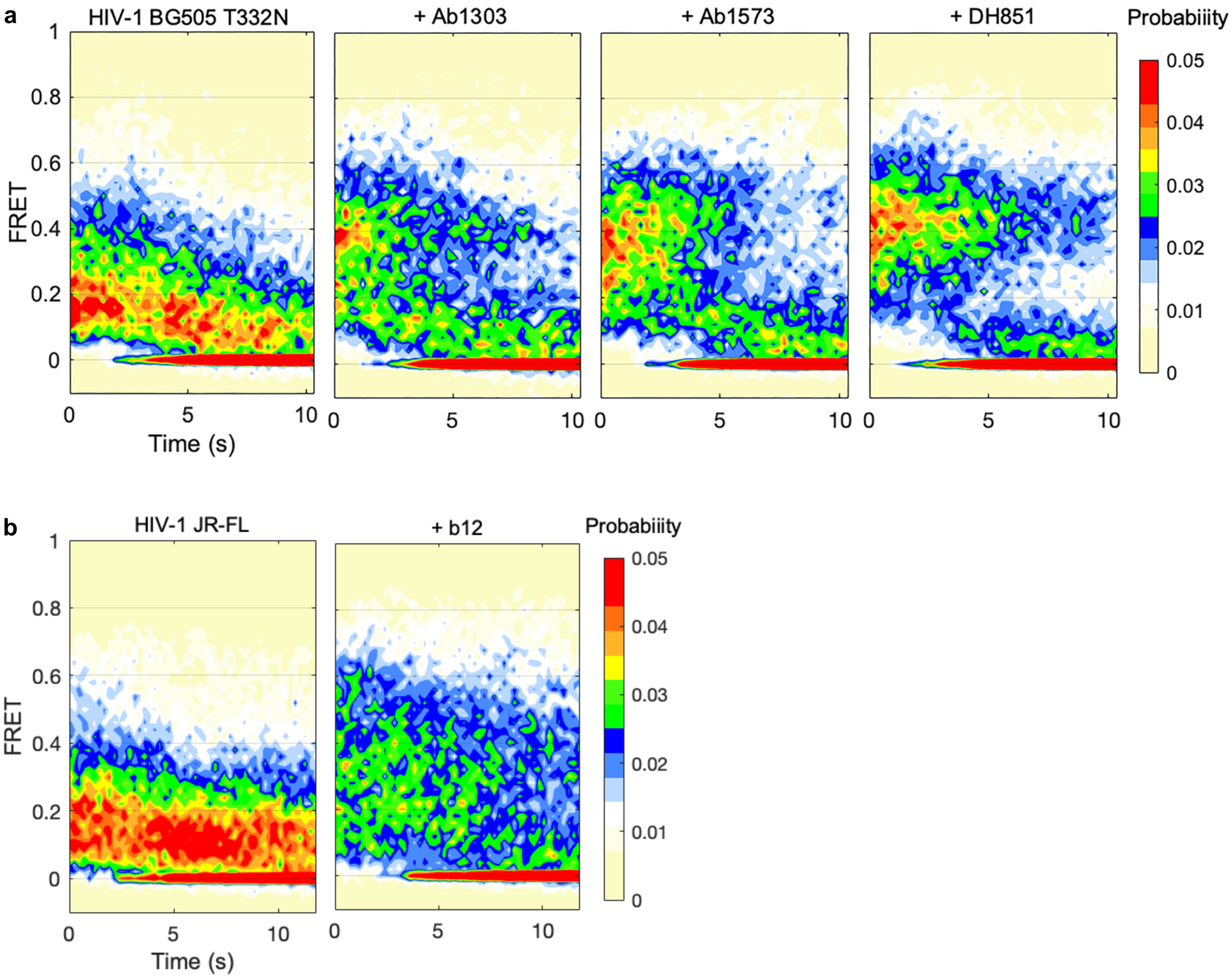


**Supplementary Fig. 2: Population contour plots of smFRET trajectories reveal peak shifting of the prevalent conformational state of Env$_{BG505}$ T332N (a) and Env$_{JR\text{-}FL}$ (b) on native virions incubated with occluded intermediate-inducing antibodies.**

Population contour plots were compiled of entire smFRET trajectories that dynamic Env trimers on intact virions undergo in the first ten seconds. The contour plots were blindly summed over time (~10 seconds) of dynamic Env trimers included in the corresponding FRET histograms (Figs. 6b – g). Fluorescently labeled Env molecules photobleached within 10 seconds contribute to the 0-FRET (baseline) population as well as baseline fluctuation variations. Population contour plots unbiasedly give overall pictures of conformational ensembles sampled by Env on native virions over time.

**a** Model fitting involves CBO (~0.3 FRET), corresponding to indicated histograms in Fig.6

| Conformational populations of Ab-incubated Env on native virions | Curve fitting $R^2$ | RMSE | PT State | PC State | CBO State |
|---|---|---|---|---|---|
| | | | μ: 0.12 ± 0.02 | μ: 0.65 ± 0.04 | μ: 0.30 ± 0.01 |
| | | | σ: 0.08 ± 0.01 | σ: 0.14 ± 0.01 | σ: 0.11 ± 0.01 |
| $Env_{BG505}$ | 0.9934 | 6.504e-4 | 46% ± 7% | 20% ± 10% | 34% ± 10% |
| $Env_{BG505}$ + Ab1303 | 0.9543 | 0.0013 | 21% ± 11% | 37% ± 16% | 42%± 19% |
| $Env_{BG505}$ + Ab1573 | 0.9710 | 9.729e-4 | 18% ± 11% | 38% ± 19% | 44% ± 22% |
| $Env_{BG505}$ + DH851 | 0.9218 | 0.0017 | 13% ± 10% | 45% ±18% | 42% ± ULM% |
| $Env_{JR-FL}$ | 0.9937 | 7.131e-4 | 55% ± 9% | 15% ± 6% | 30% ± 11% |
| $Env_{JR-FL}$ + b12 | 0.9696 | 0.0011 | 19% ± 8% | 33% ± 18% | 48% ± 19% |

**b** Model fitting involves OI (~0.4 FRET)

| Conformational populations of Ab-incubated Env on native virions | Curve fitting $R^2$ | RMSE | PT State | PC State | OI State |
|---|---|---|---|---|---|
| | | | μ: 0.12 ± 0.02 | μ: 0.65 ± 0.04 | μ: 0.4 ± 0.01 |
| | | | σ: 0.08 ± 0.01 | σ: 0.14 ± 0.01 | σ: 0.11 ± 0.01 |
| $Env_{BG505}$ | 0.9705 | 0.0013 | 54% ± 5% | 11% ± ULM% | 35% ± 8% |
| $Env_{BG505}$ + Ab1303 | 0.9904 | 6.138e-4 | 33% ± 4% | 24% ± 8% | 43% ± 9% |
| $Env_{BG505}$ + Ab1573 | 0.9813 | 8.082e-4 | 30% ± 8% | 24% ± 9% | 46% ± 9% |
| $Env_{BG505}$ + DH851 | 0.9932 | 4.903e-4 | 24% ± 5% | 27% ± 9% | 49% ± 9% |
| $Env_{JR-FL}$ | 0.9669 | 0.0016 | 62% ± 7% | 8% ± 8% | 30% ± 12% |
| $Env_{JR-FL}$ + b12 | 0.9892 | 6.460e-4 | 31% ± 8% | 22% ± 12% | 47% ± 12% |

**c** Model fitting involves both CBO (~0.3 FRET) and OI (~0.4 FRET) – overfitting and uncertainty

| Conformational populations of Ab-incubated Env on native virions | Curve fitting $R^2$ | RMSE | PT State | PC State | CBO State | OI State |
|---|---|---|---|---|---|---|
| | | | μ: 0.12 ± 0.02 | μ: 0.65 ± 0.04 | μ: 0.3 ± 0.01 | μ: 0.4 ± 0.01 |
| | | | σ: 0.08 ± 0.01 | σ: 0.14 ± 0.01 | σ: 0.06 ± 0.01 | σ: 0.06 ± 0.01 |
| $Env_{BG505}$ | 0.9967 | 4.667e-4 | 50% ± 7% | 18% ± 9% | 18% ± 15% | 14% ± 17% |
| $Env_{BG505}$ + Ab1303 | 0.9867 | 7.182e-4 | 35% ± ULM% | 31% ± ULM% | 8% ± ULM% | 26% ± ULM% |
| $Env_{BG505}$ + Ab1573 | 0.9717 | 9.917e-4 | 28% ± ULM% | 34% ± ULM% | 16% ± ULM% | 22% ± ULM% |
| $Env_{BG505}$ + DH851 | 0.9833 | 7.875e-4 | 23% ± ULM% | 38% ± ULM% | 13% ± ULM% | 26% ± ULM% |
| $Env_{JR-FL}$ | 0.9924 | 7.933e-4 | 59% ± 5% | 16% ± 12% | 12% ± 17% | 13% ± 13% |
| $Env_{JR-FL}$ + b12 | 0.9824 | 8.356e-4 | 30% ± 6% | 29% ± 13% | 15% ± 15% | 26% ± 20% |

*ULM indicates 'uncertainty larger than mean'

**d**

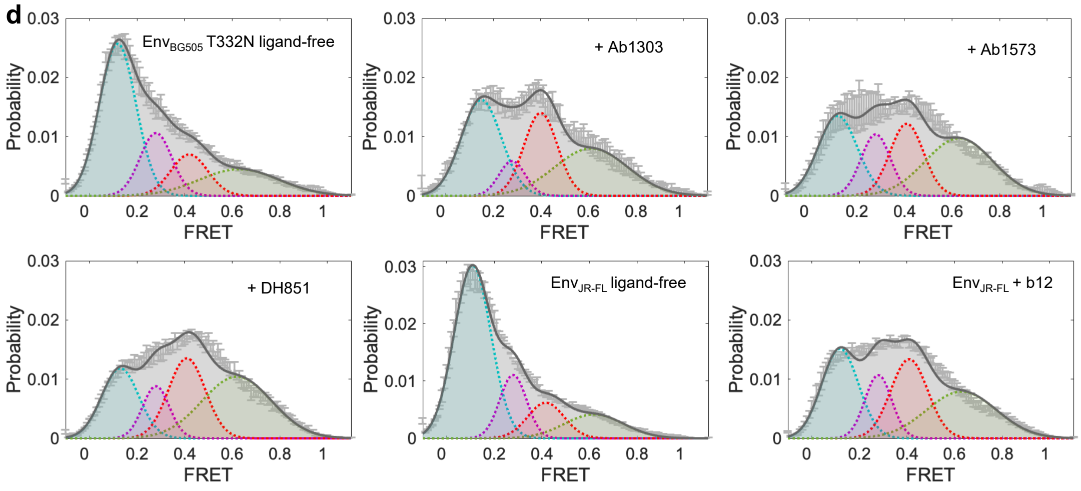


**Supplementary Fig. 3: Comparison of goodness of model fitting of FRET histograms with three and four states and fitting parameters.**

**a, b** Constrained model-fitting of FRET histograms of Ab-incubated Env on native virions into a sum of three-state Gaussian/Normal $N$ ($\mu$, $\sigma^2$) distributions that includes CBO instead of OI worsens the goodness of fitting. Parameters ($\mu$, $\sigma$) chosen for fitting were determined based on visual inspection of all trajectories that exhibit state-to-state transitions and the idealization of individual trajectories using multi-state Hidden Markov modeling. The probability of each state Env occupies was presented as mean ± s.e.m (uncertainty). ULM (s.e.m) comes from unfavorable fitting (with less than 95% confidence). $R^2$ and RMSE (Root Mean Square Deviation) evaluate the goodness of fitting. A value of $R^2$ closer to 1 and/or RMSE closer to 0 indicates a better fit/prediction. Cyan-colored $R^2$ and RMSE indicate worse fitting/prediction. Parallel comparisons of $R^2$, RMSE, and probability s.e.m obtained from model fitting involving CBO and OI, imply a better presentation of the conformational population of Ab-incubated Env using OI state than CBO state.

**c, d** Statistics and associated curve fitting of FRET histograms with a sum of four Gaussian distributions. Fitting to 4-state instead of 3-state increases the likelihood of overfitting and causes an increase in uncertainty, as implied by the elevated uncertainty of state occupancy or even ULM (generated with less than 95% confidence bounds). Uncertainty close to the level of mean was highlighted.

**a**

| Antibody-stabilized conformational states of Env on native virions | Anti-Env antibodies or ligands | | | | | | | |
|---|---|---|---|---|---|---|---|---|
| Pre-trigged (PT) | 10-1074[1] | 2G12[2] | 3BNC117[1] | PG9[2] | PG16[2] | PGT122[2] | Temsavir[1,2,3] | VRC01[2] |
| CD4-bound open (CBO) | CD4 mimetics[2] | eCD4-Ig[1] | F105[1] | sCD4$_{D1D2}$-Igαtp[1,3,4] | sCD4+17b[1,4] | | | |
| Prefusion closed (PC) | Cow-1[1] | Cow-8[1] | Cow-9[1] | Cow-10[1] | PGT151[1] | VRC34[5] | | |

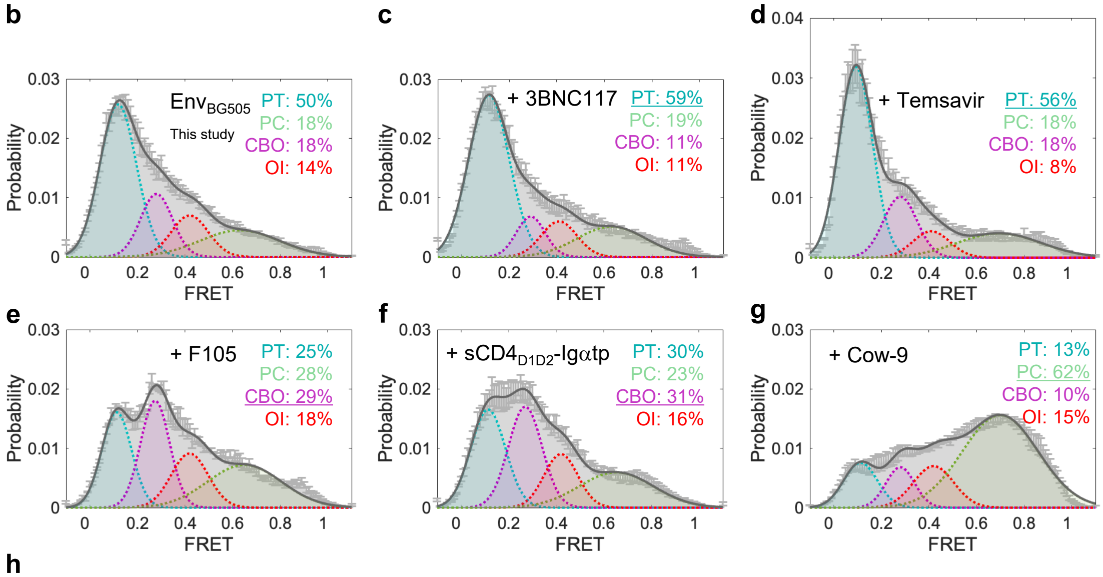


**h**

| Conformational populations of Ab-incubated Env on native virions | Curve fitting $R^2$ | RMSE | PT State | PC State | CBO State | OI State |
|---|---|---|---|---|---|---|
| | | | μ: 0.12 ± 0.02 | μ: 0.65 ± 0.04 | μ: 0.3 ± 0.01 | μ: 0.4 ± 0.01 |
| | | | σ: 0.08 ± 0.01 | σ: 0.14 ± 0.01 | σ: 0.06 ± 0.01 | σ: 0.06 ± 0.01 |
| Env$_{BG505}$ (data in this study) | 0.9967 | 4.667e-4 | 50% ± 7% | 18% ± 9% | 18% ± 15% | 14% ± 17% |
| Env$_{BG505}$ + 3BNC117 | 0.9915 | 7.481e-4 | 59% ± 6% | 19% ± 11% | 11% ± 25% | 11% ± 30% |
| Env$_{BG505}$ + Temsavir | 0.9891 | 9.352e-4 | 56% ± 3% | 18% ± 5% | 18% ± ULM% | 8% ± ULM% |
| Env$_{BG505}$ + F105 | 0.9852 | 7.777e-4 | 25% ± 4% | 28% ± ULM% | 29% ± ULM% | 18% ± 4% |
| Env$_{BG505}$ + sCD4$_{D1D2}$-Igαtp | 0.9951 | 4.810e-4 | 30% ± 2% | 23% ± 3% | 31% ± ULM% | 16% ± ULM% |
| Env$_{BG505}$ + Cow-9 | 0.9814 | 6.821e-4 | 13% ± 4% | 62% ± 7% | 10% ± 12% | 15% ± 15% |

*ULM indicates 'uncertainty larger than mean'

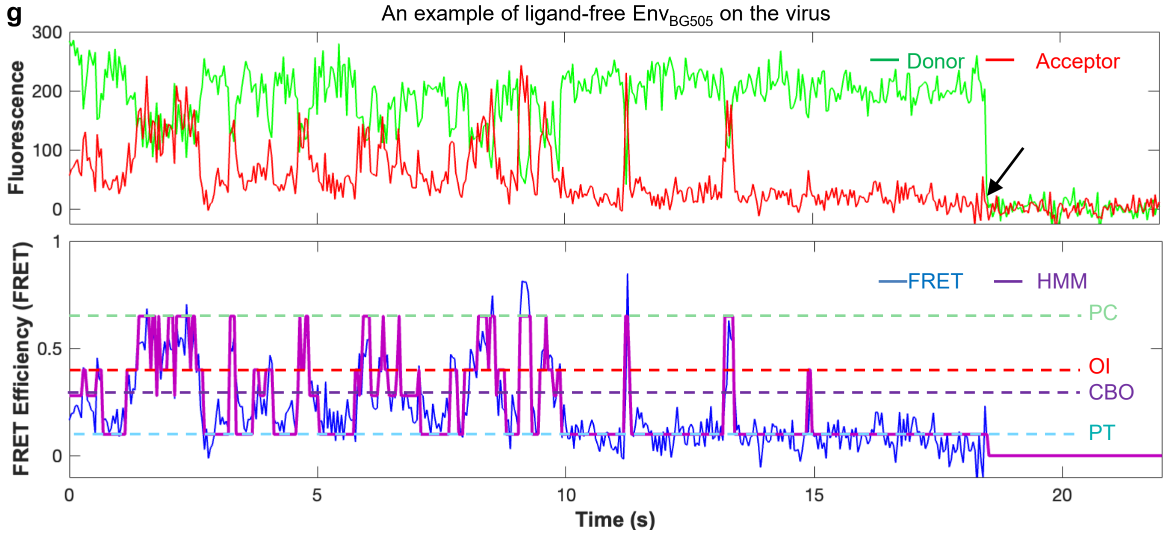


**Supplementary Fig. 4: Refitting of prior smFRET datasets with tested antibodies by 4-state Gaussian model indicates no population enrichment in the OI state.**

**a** Table listing specific stabilized states of Env by individual antibodies in previous studies (refs. 1-5).

**b – g** Example histograms describing enhancement of a prior state occupied by the ligand-free Env (**b**) by an antibody/ligand in each class, including PT-stabilizing (**c, d**), CBO-stabilizing (**e, f**) and PC-stabilizing (**g**) ligands. Histogram in b is the same as Fig.6b, whereas histograms in c – g are from published work (ref. 1). The occupancy of dominant state is underlined.

**h** Statistics and associated curve fitting of FRET histograms (**b – g**) with a sum of four Gaussian distributions, as described in Supplementary Information.

**g** Example fluorescence (donor, green; acceptor, red) trajectory and resulting FRET efficiency trajectory (FRET efficiency, blue; hidden Markov modeling – HMM idealization, magenta) of a dually labeled Env$_{BG505}$ on an intact virion. The black arrow points to the single-step photobleaching. Four FRET-populated states are indicated as color-coded dashed lines.

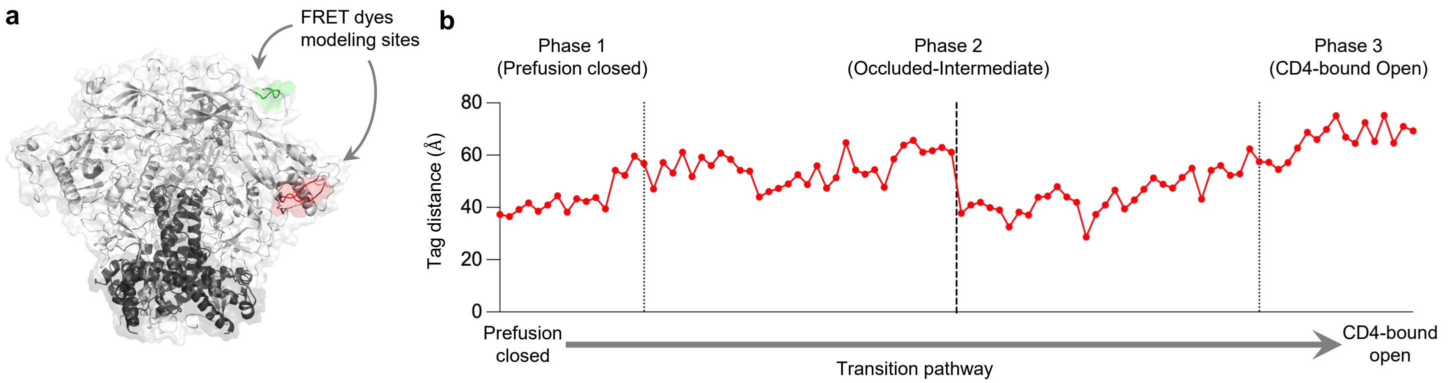


**c** FRET and tag distance of Phase 1, 2, and 3

| Phase | 1 | 2 | 3 |
|---|---|---|---|
| FRET efficiency | 0.65 | 0.4 | 0.3 |
| Tag distance (Å) | 37.3 | 28.6 – 65.7<br>(Average 50.2) | 69.2 |

**Supplementary Fig. 5: The correlation between tag distance and smFRET.**
**a** Schematic figure of modeled fluorescent donor-acceptor labels on an $Env_{BG505}$ trimer.
**b** Tag distance over the transition pathway.
**c** FRET and tag distance at each phase. The tag distances at phase 1 and 3 are from the farthest end. For phase 2, minimum to maximum value along with average distance are shown.

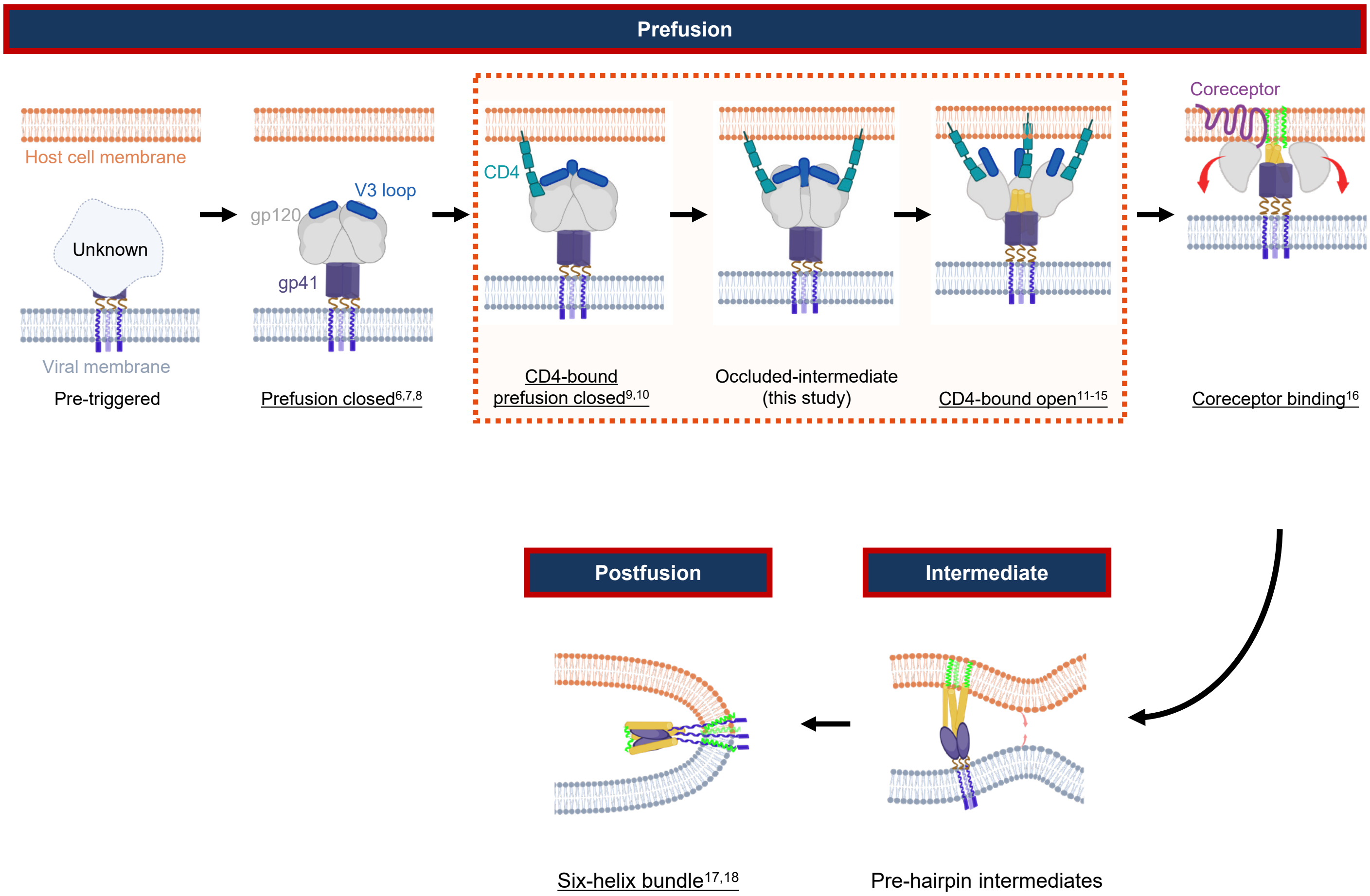


**Supplementary Fig. 6: Schematic figure depicting the scope of current study in orange box over HIV-1 Env fusion process.**
Note that for logistical reasons, dynamics were performed with three CD4s throughout the trajectory. Conformations with underline labels have determined residue-level structures.
